## Supplementary material for "Unistrand piRNA clusters are an evolutionarily conserved mechanism to suppress endogenous retroviruses across the *Drosophila* genus": Figure S1-S26 and list of supplementary tables

**Supplementary Figures**

**Fig. S1:** Identification of *flam* within species of the *melanogaster* subgroup
**Fig. S2:** Identification of *flam* in species outside of the *melanogaster* subgroup
**Fig. S3:** RepeatMasker underestimates interspersed repeat content in distant species
**Fig. S4:** Identification of *flam* clusters using genome-wide LTR content
**Fig. S5:** Identification of *flam* in species outside of the *melanogaster* subgroup
**Fig. S6:** Identification of *flam*-like clusters using genome-wide LTR content
**Fig. S7:** *Flamlike3* across the *obscura* group
**Fig. S8:** Examples of difficult cases in the identification of *flam*-syntenic clusters
**Fig. S9:** Overview of additional *flamlike3*- and *flamlike5*-syntenic regions
**Fig. S10:** *Flamlike5* across the *obscura* group
**Fig. S11:** *Flamlike5*-syntenic regions across *pseudoobscura* subgroup
**Fig. S12:** Conservation of ATAC-seq peaks in the *flam* promoter region of the *melanogaster* subgroup
**Fig. S13:** ATAC-seq peaks in the promoter region of *flamlike3* and *flamlike1*
**Fig. S14:** *Flam*-like and unistrand piRNA clusters are somatically expressed
**Fig. S15:** Somatic piRNA clusters are expressed from one strand
**Fig. S16:** Chromosomal location of somatic piRNA clusters with *flam*-like properties
**Fig. S17:** Overview of *de novo* transposon consensus sequence construction
**Fig. S18:** Repeat content of *Drosophila* species
**Fig. S19:** *De novo* TE libraries enable detailed cluster analyses
**Fig. S20:** Overview of identified subfamilies per TE family across *Drosophila* species
**Fig. S21:** Overview of genomic copies per TE family across *Drosophila* species
**Fig. S22:** Overview of previously described TE subfamilies across *Drosophila* species
**Fig. S23:** Alignment of sRNA-seq across *Drosophila* species
**Fig. S24:** Alignment of RNA-seq across *Drosophila* species
**Fig. S25:** *Flam*-like clusters display somatic expression and reduced ping-pong signature
**Fig. S26:** Conservation of *fs(1)Yb* across *Drosophila* species

**Supplementary Tables**

**Table S1:** Coordinates for all unistrand flam-like piRNA cluster candidates identified in this study.
**Table S2:** Prediction of major de-novo piRNA clusters using somatic and total sRNA-seq data across all species sequenced in this study.
**Table S3:** Genomic coverage for each transposon family across all 193 assembles.

**Table S4:** Number of subfamilies annotated as each transposon family across all 193 assembles.
**Table S5:** Genomic copies per transposon family across all 193 assembles.
**Table S6:** Detection of 155 previously described transposon subfamilies across the curated de-novo transposon libraries.
**Table S7:** List of all 119 species and 193 assemblies included in this study and abbreviations used.
**Table S8:** Information about the fly species included in this study.
**Table S9:** List of all sRNA-seq libraries analysed in this study and their alignment metrics.
**Table S10:** List of all RNA-seq libraries analysed in this study and their alignment metrics.
**Table S11:** List of all ATAC-seq libraries analysed in this study and their alignment metrics.

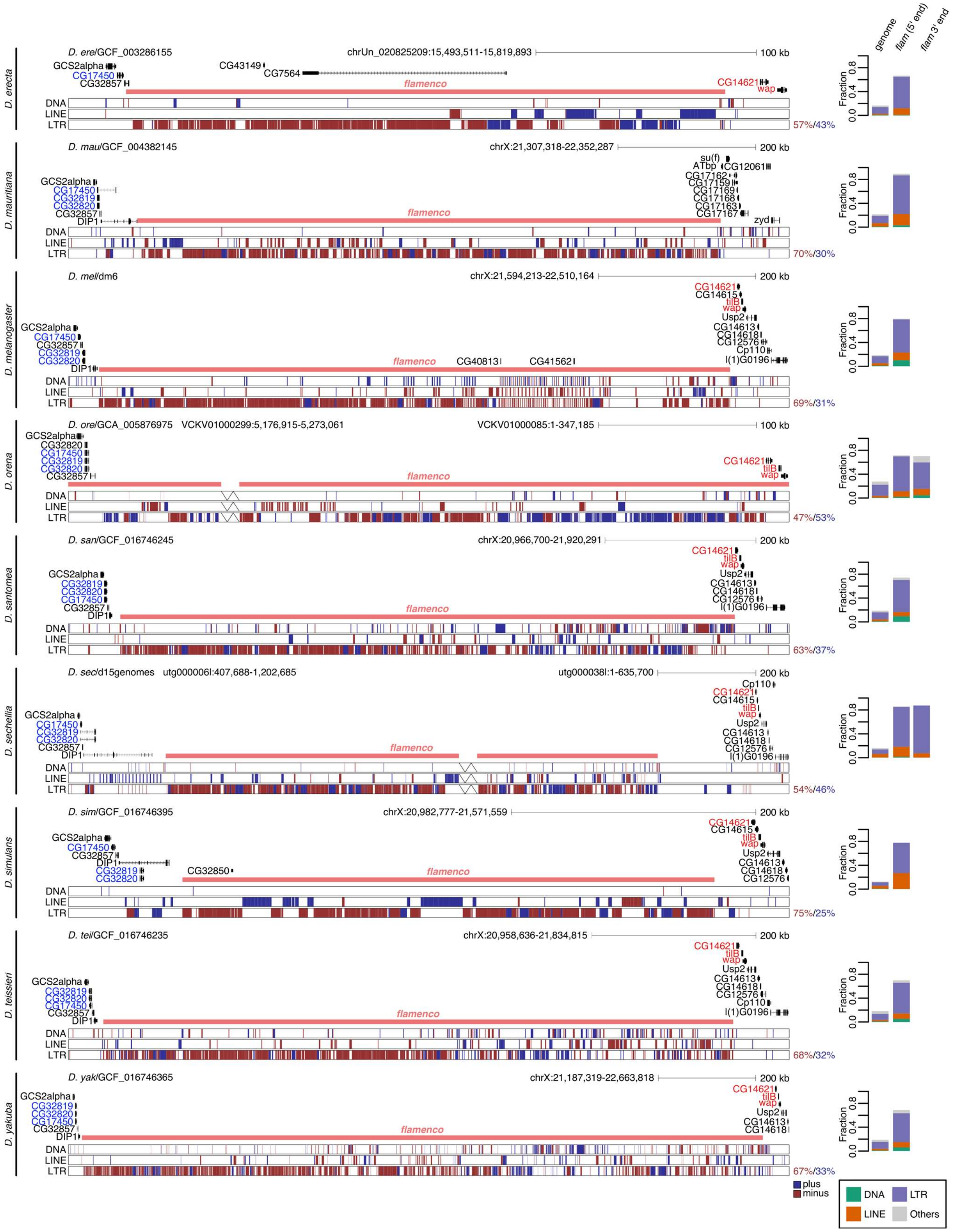


**Fig. S1: Identification of *flam* within species of the *melanogaster* subgroup**

Genome browser tracks showing *flam* region for species within the *melanogaster* subgroup. Repeat content was annotated using RepeatMasker. Gene annotations show *D. melanogaster* transcripts mapped onto the target genome. Assembly breakpoints are indicated by a break line symbol. Percentage to the right indicates LTR content per strand. Bar graph shows fraction of annotated TEs in the genome and the *flam*-syntenic region, respectively, split in 5' and 3' end if there was an assembly breakpoint.

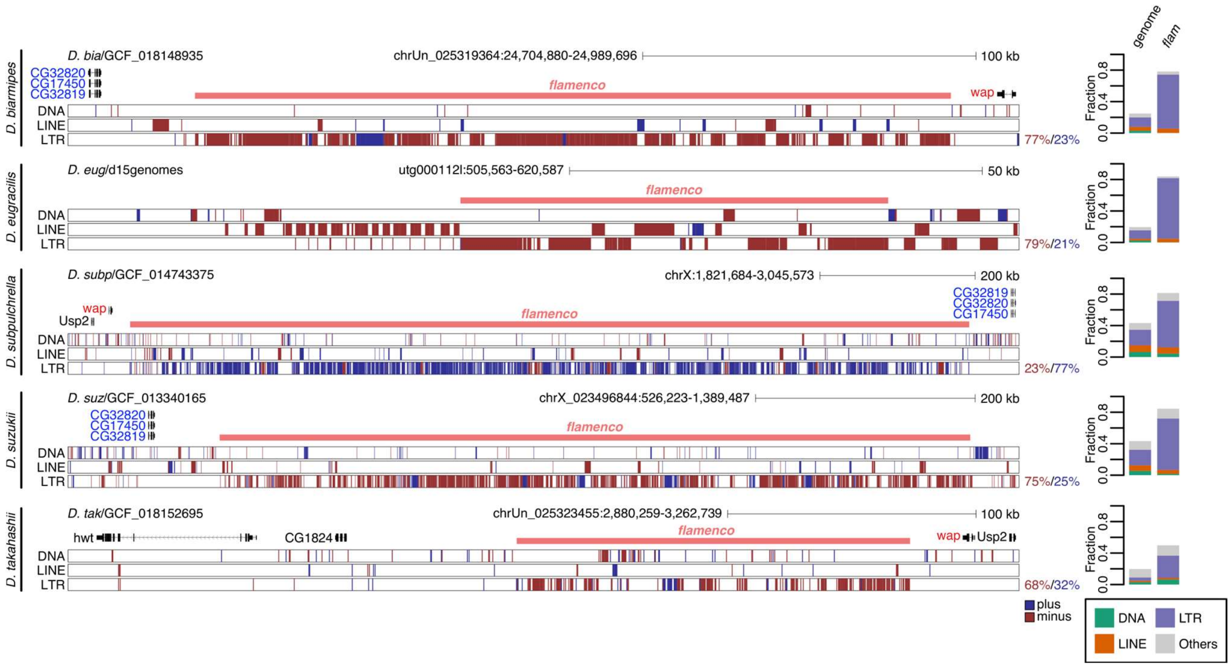


**Fig. S2: Identification of *flam* in species outside of the *melanogaster* subgroup**

Genome browser tracks showing *flam* region for species outside of the *melanogaster* subgroup. Repeat content was annotated using RepeatMasker. Gene annotations show *D. melanogaster* transcripts mapped onto the target genome. Percentage to the right indicates LTR content per strand. Bar graph shows fraction of annotated TEs in the genome and the *flam*-syntenic region, respectively.

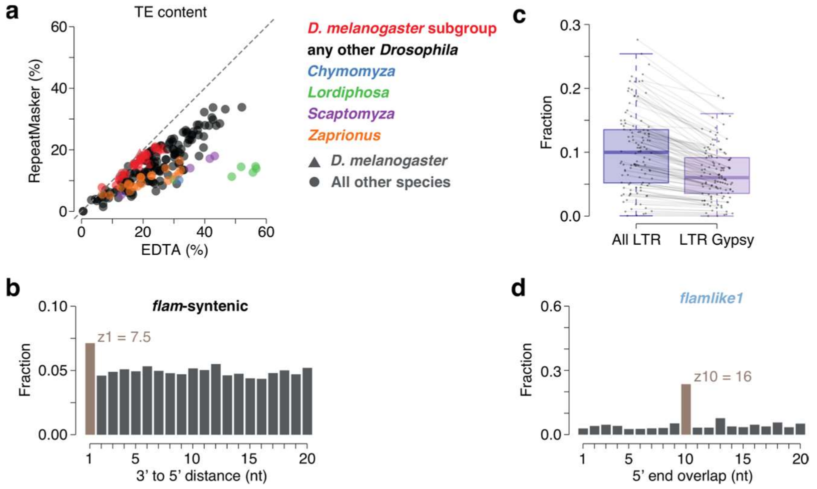


**Fig. S3: RepeatMasker underestimates interspersed repeat content in distant species**

**a**, Genomic interspersed repeat content per genome assembly predicted using RepeatMasker and EDTA, respectively. Genome assemblies are coloured per genus. **b**, Phasing signature (3’ end to 5’ end distance) for piRNA pairs mapping onto the *flam*-syntenic region. **c**, Fraction of LTR TEs and LTR *Gypsy* TEs for each species (n=119). Species with multiple genome assemblies are represented by their mean. Lines connect each species between the two box plots. **d**, Ping-pong signature for piRNA pairs mapping onto the *flamlike1* region.

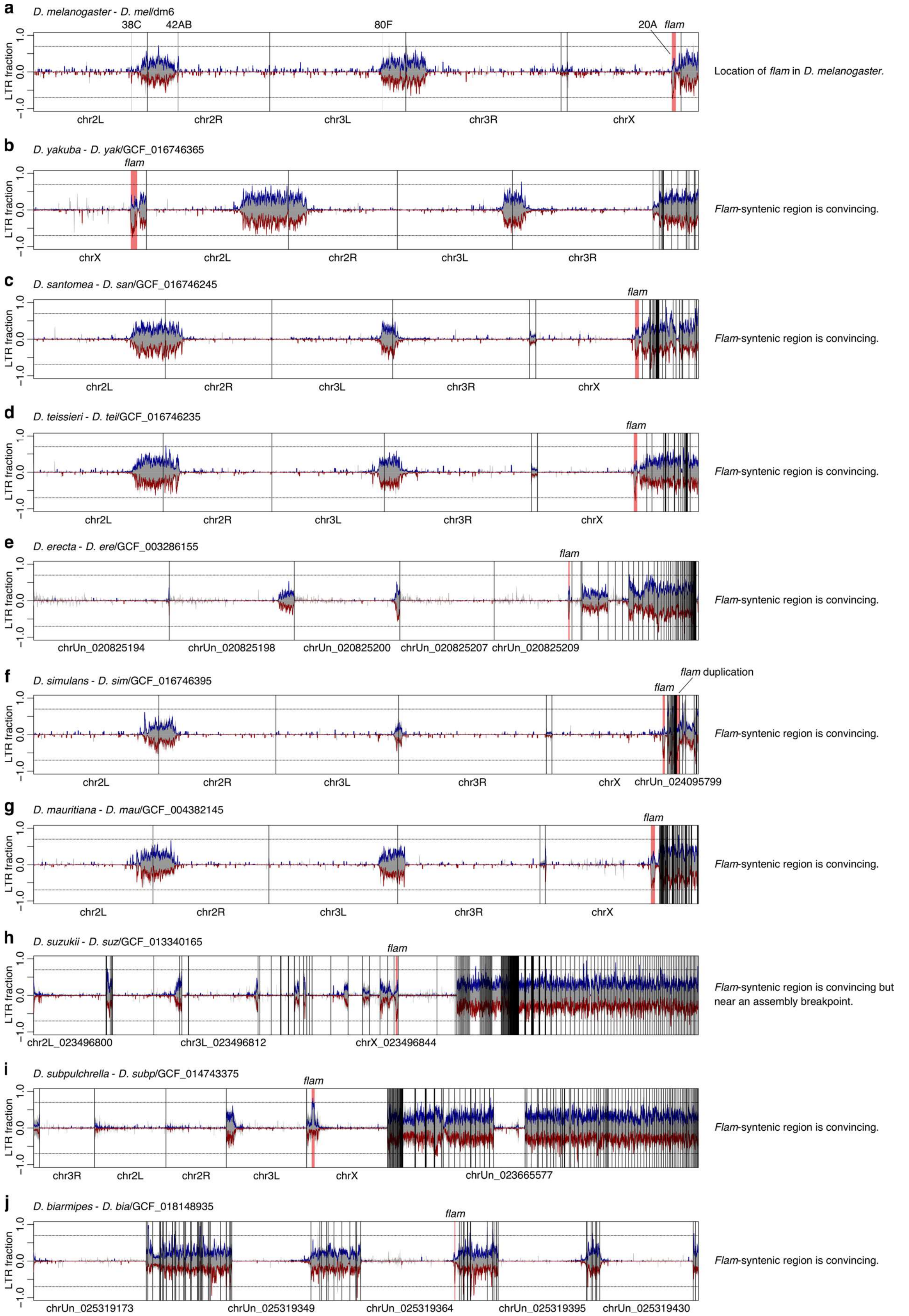


**Fig. S4: Identification of *flam* clusters using genome-wide LTR content**

**a-j**, The genome was scanned for *flam*-like clusters based on genomic LTR content across 100 kb windows. Location of (a) *flam* and other major clusters in *D. melanogaster* and (b-j) a *flam*-syntenic region across nine other species. Predicted LTR content (blue, plus strand; red, minus strand) and total repeat content (grey) is shown across the whole genome (100 kb windows). Cluster loci are indicated at the top and selected contig/chromosome names are indicated at the bottom.

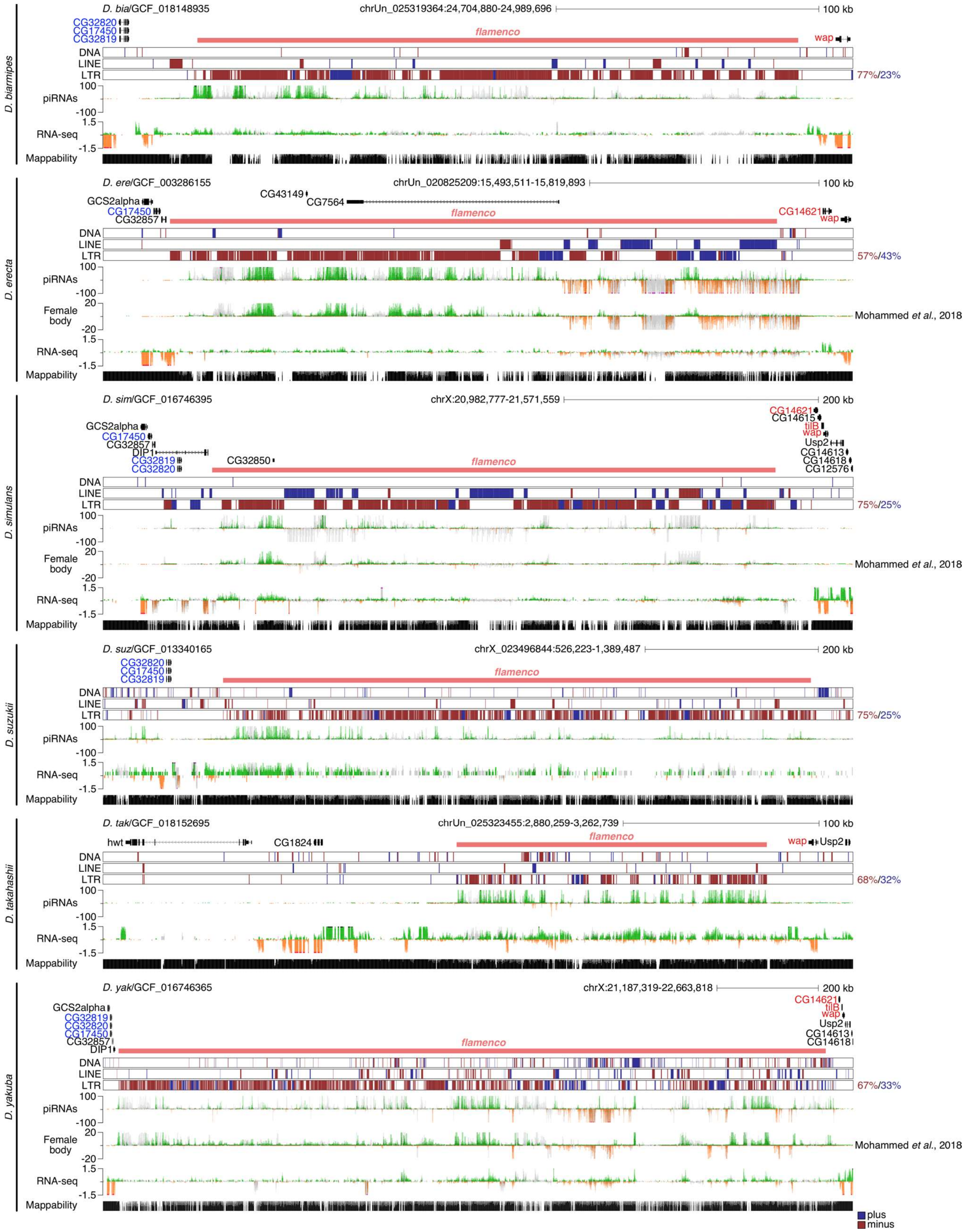


**Fig. S5: Identification of *flam* in species outside of the *melanogaster* subgroup**

Genome browser tracks showing *flam* region for indicated species. Repeat content was annotated using EDTA. Gene annotations show *D. melanogaster* transcripts mapped onto the target genome. Percentage to the right indicates LTR content per strand. Uniquely mapping piRNAs (cpm) and total RNA expression (ln(cpm+1)) and mappability is shown where available. Sequencing data is shown in green or orange for uniquely mapped reads, and grey for multi-mapping reads. Publicly available sRNA data is indicated [60].

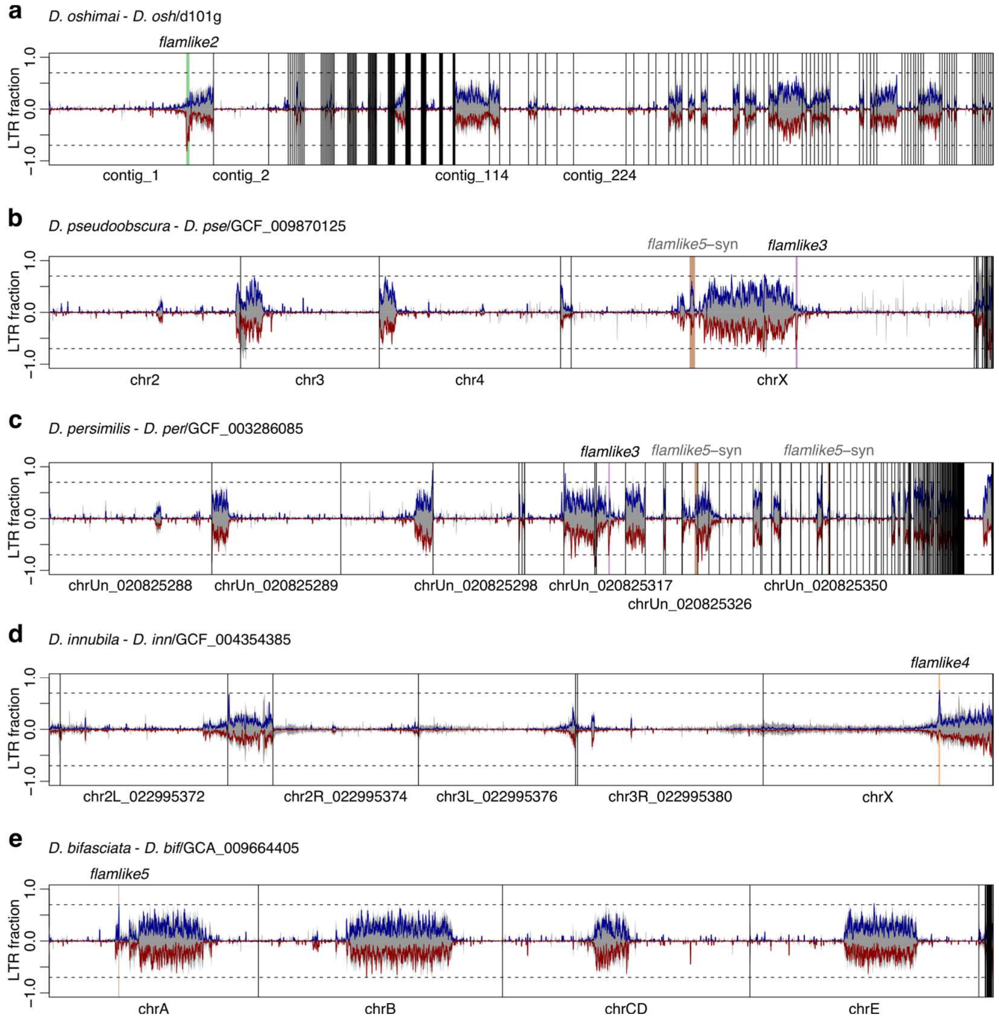


**Fig. S6: Identification of *flam*-like clusters using genome-wide LTR content**

**a-e**, The genome was scanned for *flam*-like clusters based on genomic LTR content across 100 kb windows. Predicted LTR content (blue, plus strand; red, minus strand) and total repeat content (grey) is shown across the whole genome (100 kb windows). Location of (a) *flamlike2*, (b-c) *flamlike3*, (d) *flamlike4*, and (e) *flamlike5* loci are indicated at the top and selected contig/chromosome names are indicated at the bottom. chrA-E refer to the Müller element nomenclature, where chrA corresponds to the X chromosome [73].

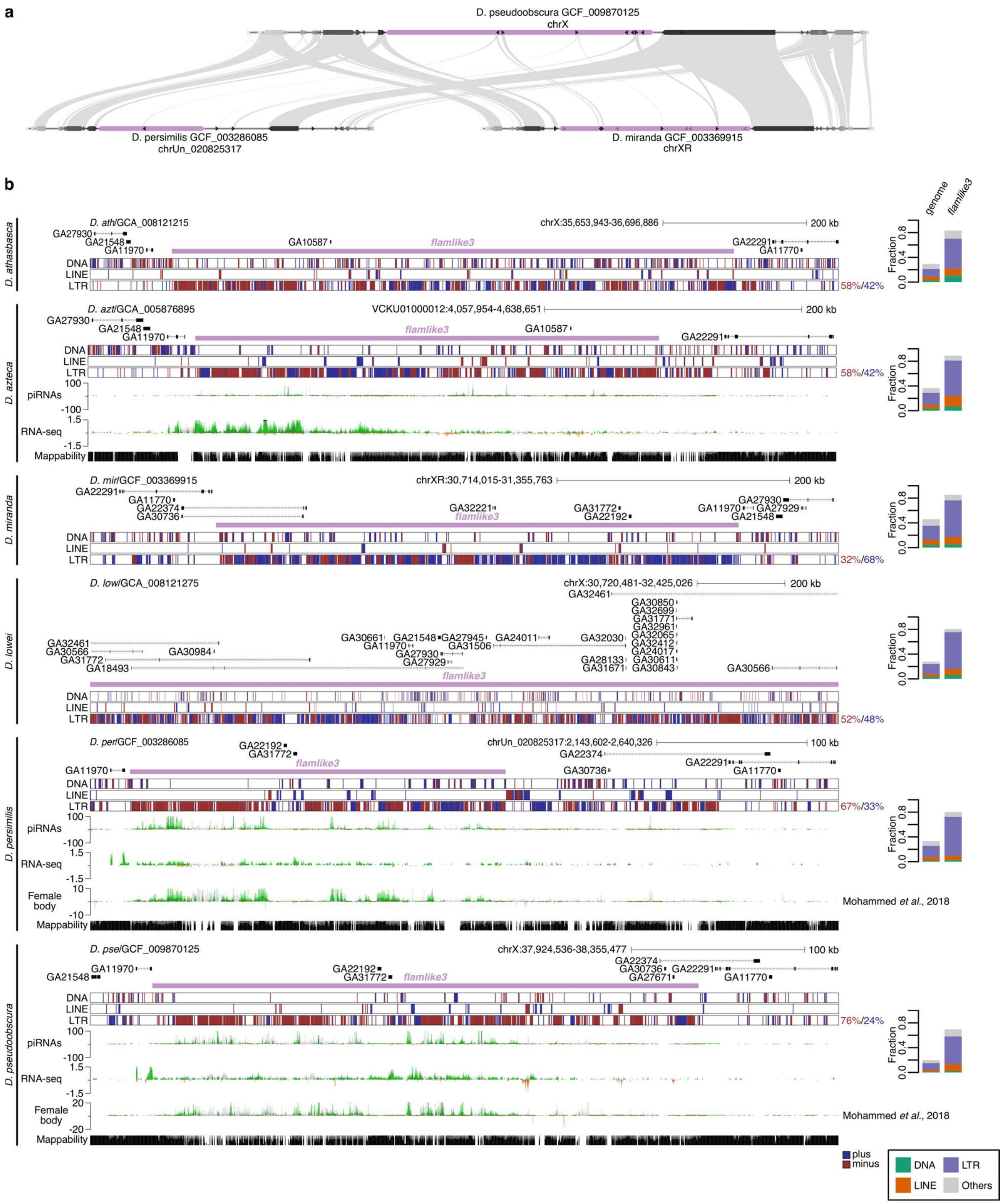

**Fig. S7: *Flamlike3* across the *obscura* group**

**a**, MCScan plot showing synteny between three species in a region containing *flamlike3*. **b**, Genome browser tracks showing *flamlike3* region for species within the *obscura* group. Repeat content was annotated using EDTA. Gene annotations show *D. pseudoobscura* transcripts mapped onto the target genome. Percentage to the right indicates LTR content per strand. Bar graph shows fraction of annotated TEs in the complete genome and the *flamlike3* region. Uniquely mapping piRNAs (cpm) and total RNA expression (ln(cpm+1)) and mappability is shown where available. Sequencing data is shown in green or orange for uniquely mapped reads, and grey for multi-mapping reads. Publicly available sRNA data is indicated [60].

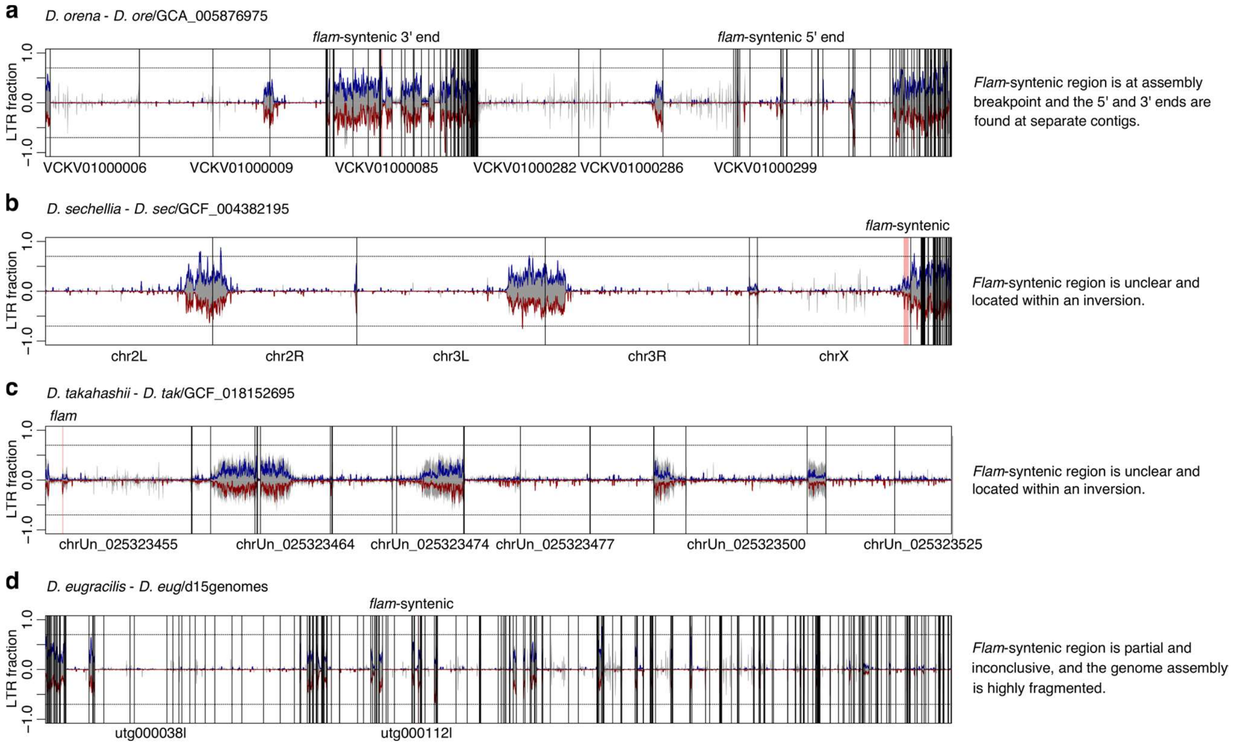


**Fig. S8: Examples of difficult cases in the identification of *flam*-syntenic clusters**

**a-d**, The genome was scanned for *flam*-like clusters based on genomic LTR content across 100 kb windows. Predicted LTR content (blue, plus strand; red, minus strand) and total repeat content (grey) is shown across the whole genome (100 kb windows). Location of a *flam*-syntenic region across four species that were not identified using the initial genome-wide scanning approach. Cluster locations based on gene synteny are indicated at the top and selected contig/chromosome names are indicated at the bottom. Due to difficulties in defining a unistranded region, only *D. takahashii* (c) was considered to have a *flam* cluster and the others (a, b, d) were considered inconclusive and potentially dual-strand *flam*-syntenic regions.

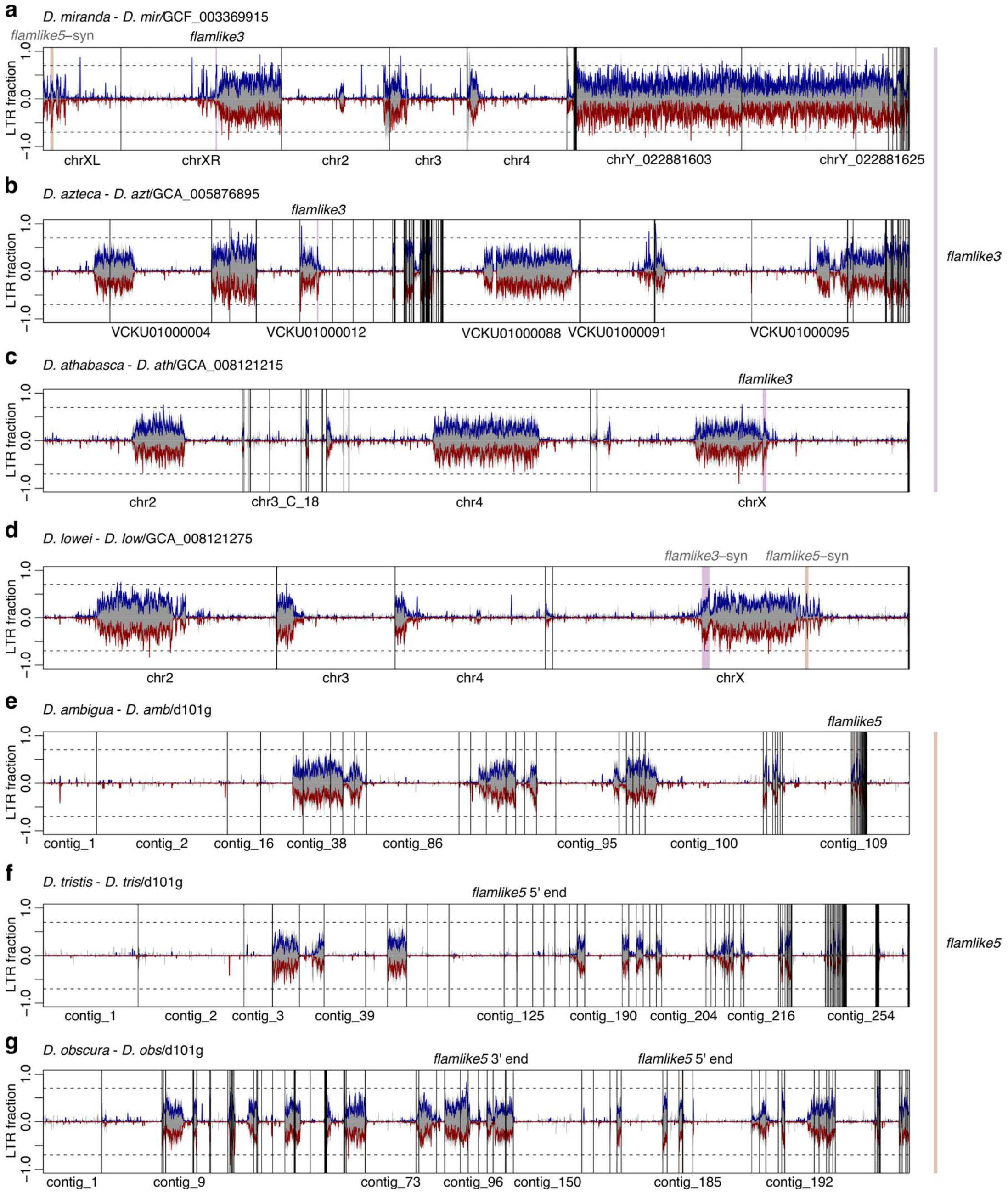


**Fig. S9: Overview of additional *flamlike3*- and *flamlike5*-syntenic regions**

**a-g**, The genome was scanned for *flam*-like clusters based on genomic LTR content across 100 kb windows. Predicted LTR content (blue, plus strand; red, minus strand) and total repeat content (grey) is shown across the whole genome (100 kb windows). Location of (a-c) *flamlike3*, (d) neither, and (e-g) *flamlike5* across *obscura* group species that were not identified using the initial genome-wide scanning approach. Cluster locations based on gene synteny are indicated at the top and selected contig/chromosome names are indicated at the bottom.

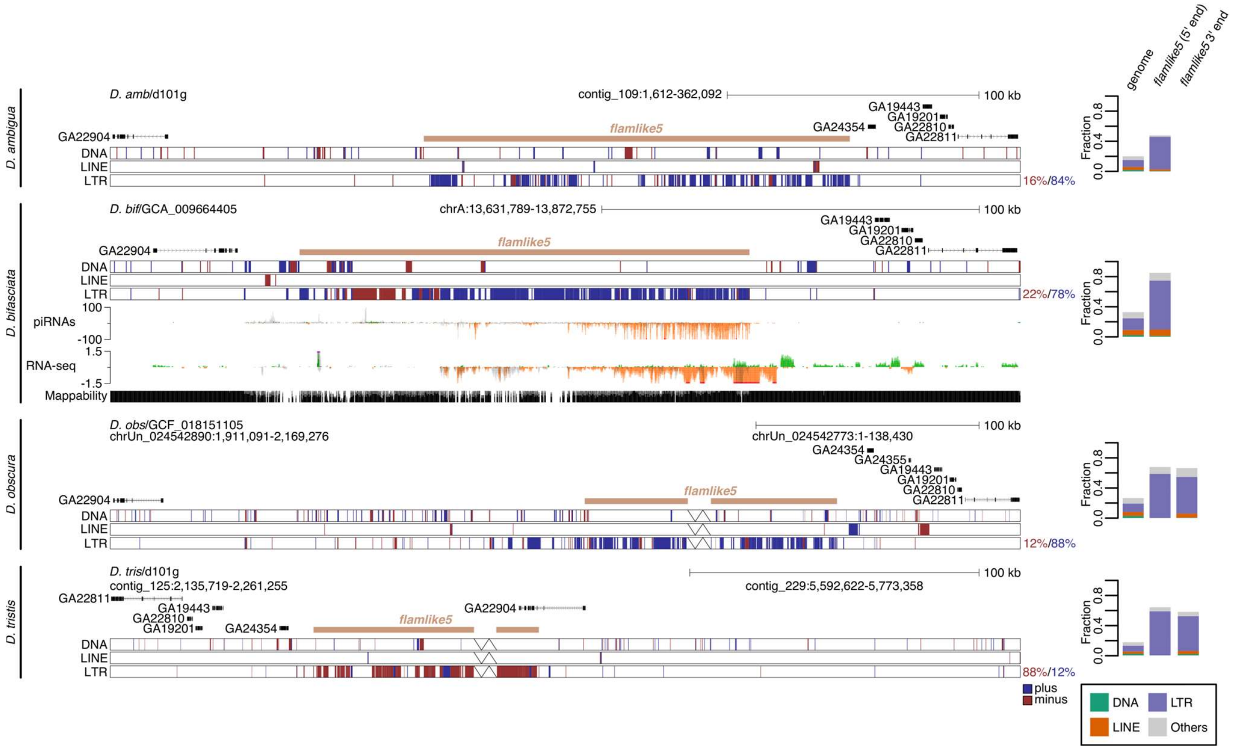


**Fig. S10:** ***Flamlike5*** **across the** ***obscura*** **group**

Genome browser tracks showing *flamlike5* region for species within the *obscura* group. Repeat content was annotated using EDTA. Gene annotations show *D. pseudoobscura* transcripts mapped onto the target genome. Assembly breakpoints are indicated by a break line symbol. Percentage to the right indicates LTR content per strand. Bar graph shows fraction of annotated TEs in the genome and the *flamlike5* region, respectively, split in 5' and 3' end if there was an assembly breakpoint. The 3' end contigs have been reverse complemented for *D. obscura* and *D. tristis*. Uniquely mapping piRNAs (cpm) and total RNA expression (ln(cpm+1)) and mappability is shown where available. Sequencing data is shown in green or orange for uniquely mapped reads, and grey for multi-mapping reads.

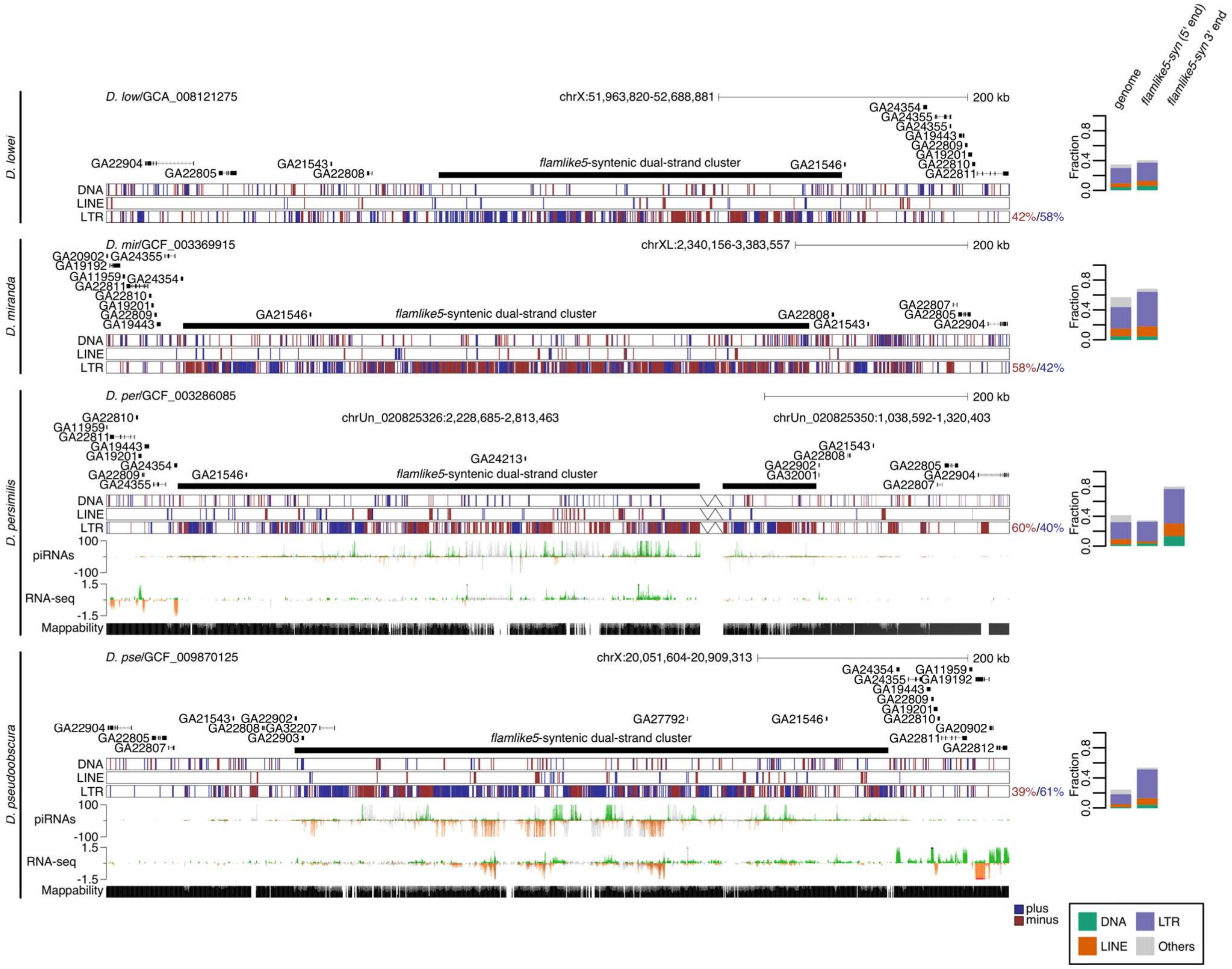


**Fig. S11: *Flamlike5*-syntenic regions across *pseudoobscura* subgroup**

Genome browser tracks showing *flamlike5*-syntenic regions for species within the *pseudoobscura* group. Repeat content was annotated using EDTA. Gene annotations show *D. pseudoobscura* transcripts mapped onto the target genome. Assembly breakpoints are indicated by a break line symbol. Percentage to the right indicates LTR content per strand. Bar graph shows fraction of annotated TEs in the complete genome and the *flamlike5* region, split in 5' and 3' end if there was an assembly breakpoint. Uniquely mapping piRNAs (cpm) and total RNA expression (ln(cpm+1)) and mappability is shown where available. Sequencing data is shown in green or orange for uniquely mapped reads, and grey for multi-mapping reads.

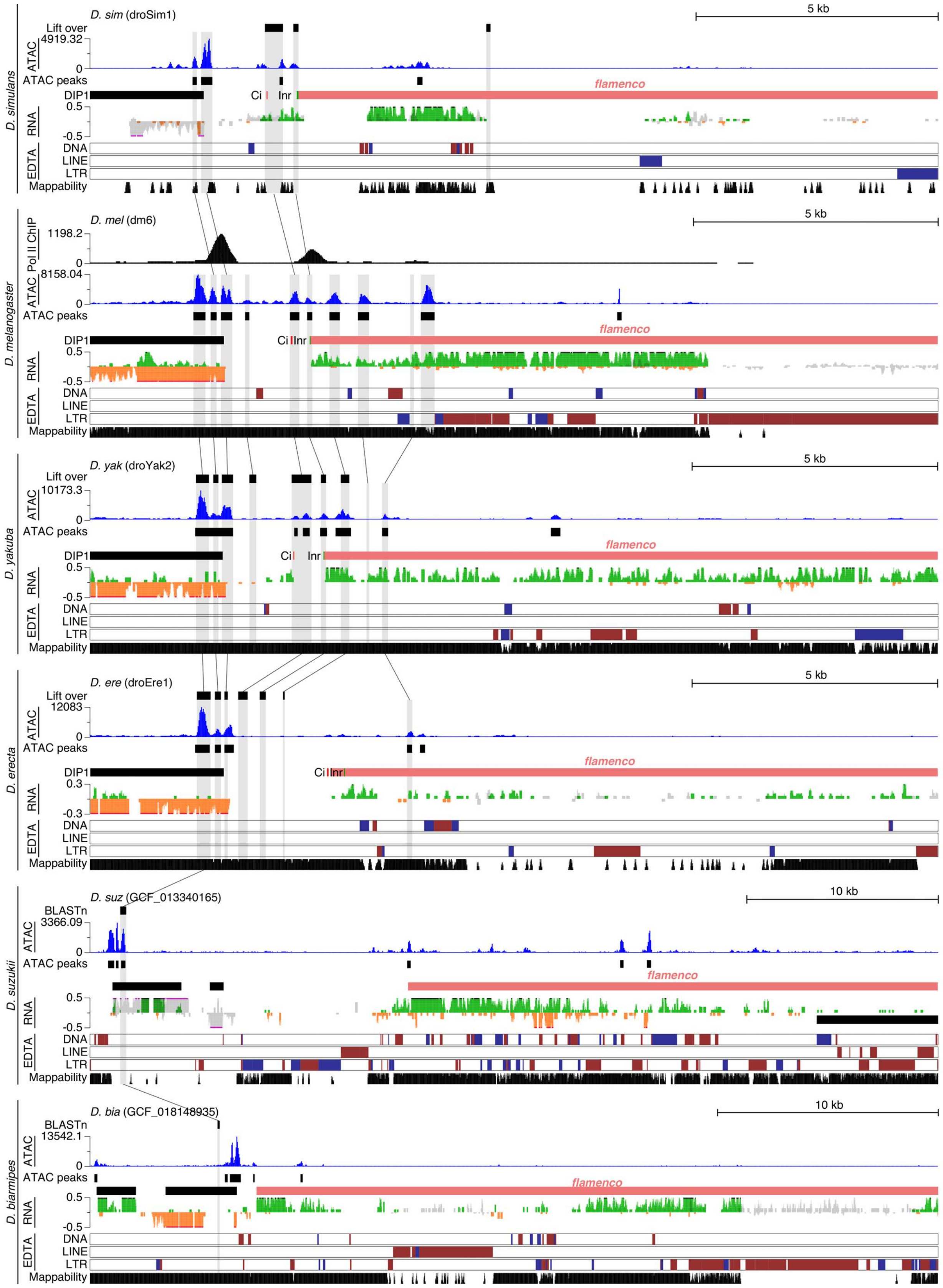

D. simulans
D. sim (droSim1)
5 kb
Lift over
ATAC
4919.32
0
ATAC peaks
DIP1
Ci
Inr
flamenco
RNA
0.5
-0.5
EDTA
DNA
LINE
LTR
Mappability
D. melanogaster
D. mel (dm6)
5 kb
Pol II ChIP
1198.2
0
ATAC
8158.04
0
ATAC peaks
DIP1
Ci
Inr
flamenco
RNA
0.5
-0.5
EDTA
DNA
LINE
LTR
Mappability
D. yakuba
D. yak (droYak2)
5 kb
Lift over
ATAC
10173.3
0
ATAC peaks
DIP1
Ci
Inr
flamenco
RNA
0.5
-0.5
EDTA
DNA
LINE
LTR
Mappability
D. erecta
D. ere (droEre1)
5 kb
Lift over
ATAC
12083
0
ATAC peaks
DIP1
Ci
Inr
flamenco
RNA
0.3
-0.3
EDTA
DNA
LINE
LTR
Mappability
D. suzukii
D. suz (GCF_013340165)
10 kb
BLASTn
ATAC
3366.09
0
ATAC peaks
flamenco
RNA
0.5
-0.5
EDTA
DNA
LINE
LTR
Mappability
D. biarmipes
D. bia (GCF_018148935)
10 kb
BLASTn
ATAC
13542.1
0
ATAC peaks
flamenco
RNA
0.5
-0.5
EDTA
DNA
LINE
LTR
Mappability

**Fig S12: Conservation of ATAC-seq peaks in the *flam* promoter region of the *melanogaster* subgroup**

**a-f**, Genome browser tracks showing *flam* region for indicated species. Uniquely mapping ATAC-seq reads (rpkm) are shown, highlighting the called ATAC-seq peaks (black bars). Peak-orthologous regions (LiftOver or BLAST) are highlighted in light grey. The position of the Ci motif and Inr element are indicated in red and green, respectively. Total RNA expression (cpm) is shown in green (forward strand) or orange (reverse strand) for uniquely mapped reads, and grey for multi-mapping reads. Repeat content was annotated using EDTA. Gene annotations show *D. melanogaster* transcripts mapped onto the target genome. Genome mappability tracks are shown for each species. Abbreviations: Ci, cubitus interruptus; Inr, Initiator element.

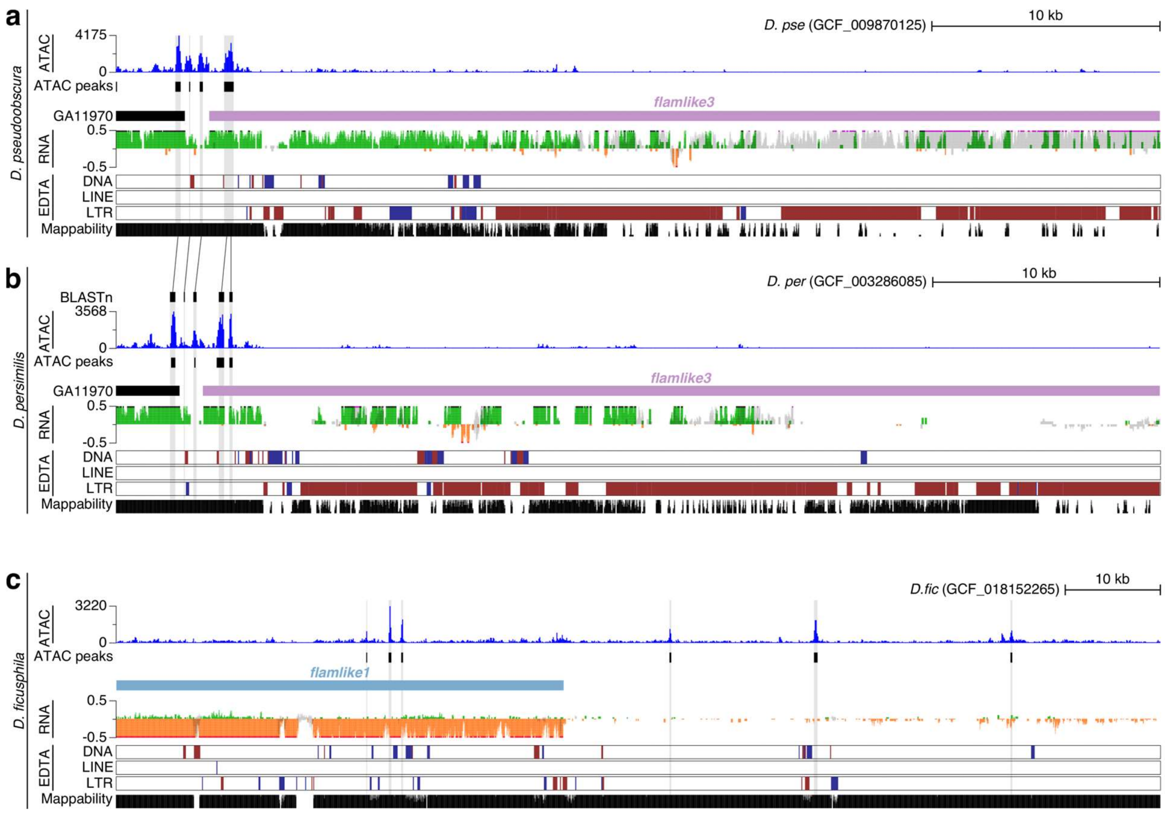


**Fig S13: ATAC-seq peaks in the promoter region of *flamlike3* and *flamlike1***

**a-b**, Genome browser tracks showing ATAC-seq peaks in the promoter of *flamlike3* for *D. pseudoobscura* (a) and *D. persimilis* (b). Uniquely mapping ATAC-seq reads (rpkm) are shown, highlighting the called ATAC-seq peaks (black bars). Peak-orthologous regions (LiftOver or BLAST) are highlighted in light grey. Total RNA expression (cpm) is shown in green (forward strand) or orange (reverse strand) for uniquely mapped reads, and grey for multi-mapping reads. Repeat content was annotated using EDTA. Gene annotations show *D. pseudoobscura* transcripts mapped onto the target genome. Genome mappability tracks are shown for each species. **c**. same as (a-b) but showing the promoter region of *flamlike1* in *D. ficusphila*. Please note that the cluster is located on the reverse strand.

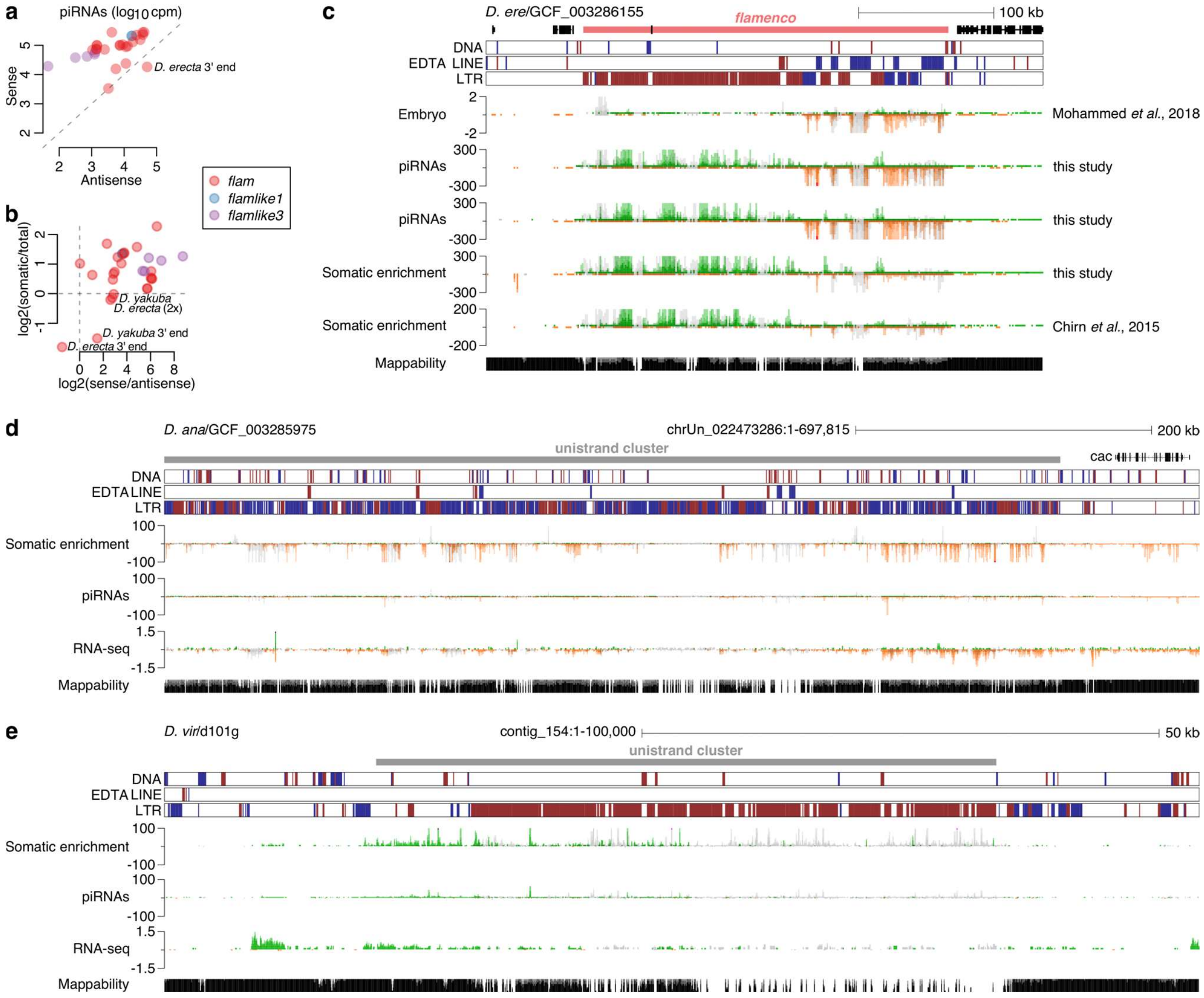


**Fig. S14: *Flam*-like and unistrand piRNA clusters are somatically expressed**

**a**, piRNAs mapping to piRNA clusters in sense or antisense orientation across all sequenced species (n=10). All clusters across all genome assemblies are shown. Some clusters have their 3' and 5' ends on different contigs. Clusters that deviate from the major strand bias are indicated. **b**, Scatterplot of piRNA strand bias against piRNA soma enrichment across the indicated clusters (n=10 species). All clusters across all genome assemblies are shown. Some clusters have their 3' and 5' ends on different contigs. Clusters that lack soma enrichment are indicated. **c**, Genome browser tracks of *flam* in *D. erecta*. *Flam* is located on the plus strand with its 5' end to the left and 3' end to the right. The sRNA-seq tracks are sorted according to germline to somatic expression patterns. Transposon annotations are shown in red (minus strand) or blue (plus strand). Sequencing data is shown in green or orange for uniquely mapped reads, and grey for multi-mapping reads. piRNA abundance is shown as counts per million. Publicly available data is indicated [26,60]. **d**, Genome browser tracks of a somatic piRNA cluster in *D. ananassae*. Transposon annotations are shown in red (minus strand) or blue (plus strand). Sequencing data is shown in green or orange for uniquely mapped reads, and grey for multi-mapping reads. piRNA abundance is shown as counts per million. RNA-seq is shown as ln(cpm+1). **e**, As in (d) but showing a somatic piRNA cluster in *D. virilis*.

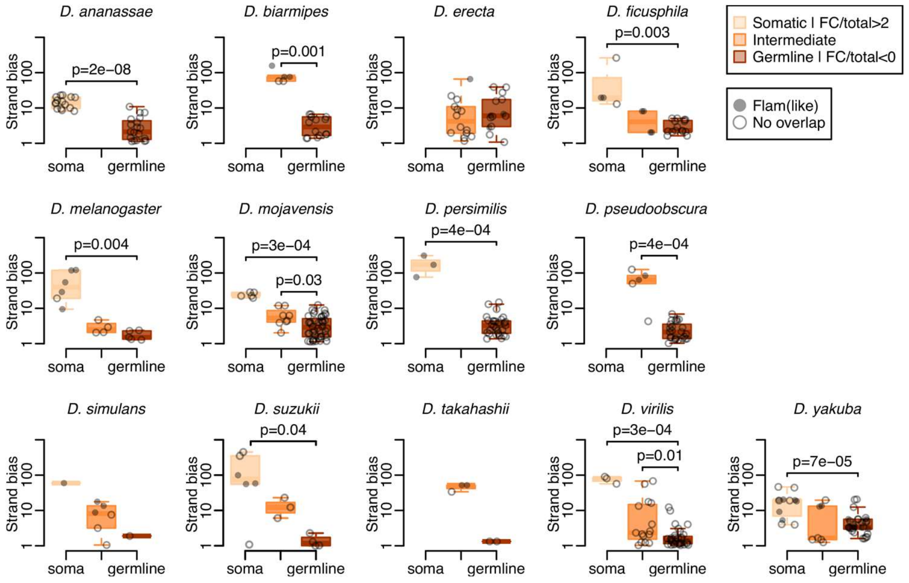


**Fig. S15: Somatic piRNA clusters are expressed from one strand**

Analysis across *de novo* identified large piRNA clusters (proTRAC, >35kb; see Methods). Clusters were classified as somatic, intermediate or germline based on the follicle cell versus total ovary piRNA ratio. Overlap to previously identified clusters (filled circle) or no overlap (empty circle) is indicated. Strand bias ($\log_{10}$ scale) is shown across each category. Germline strand bias was compared to intermediate or somatic using a two-sided Wilcoxon's rank sum test. The same cluster may be represented multiple times if it was identified in multiple genome assemblies. Raw data and cluster coordinates are available in Table S2. Boxplots show median (central line), interquartile range (IQR, box), and minimum and maximum values (whiskers, at most 1.5*IQR).

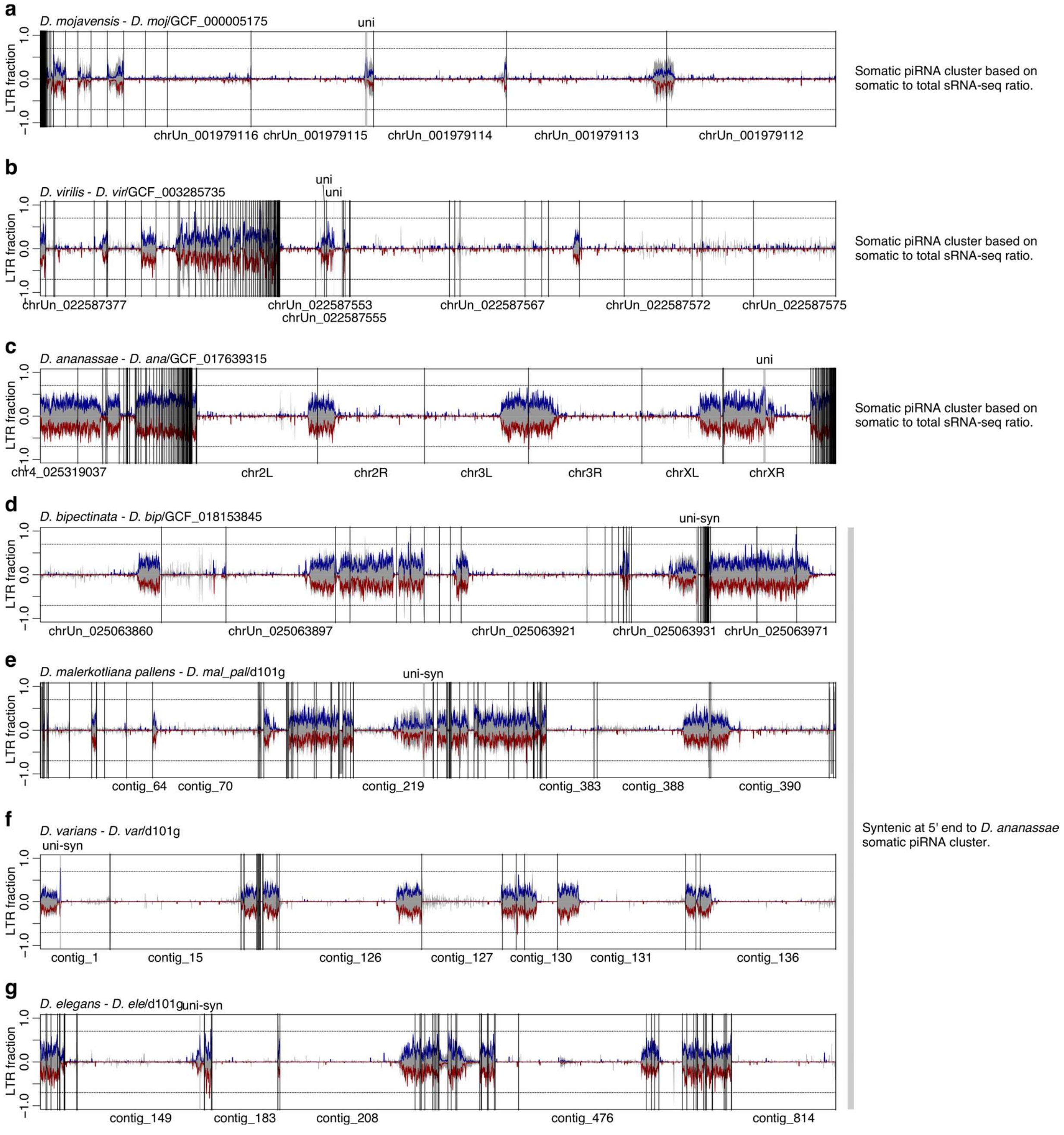


**Fig. S16: Chromosomal location of somatic piRNA clusters with *flam*-like properties**
**a-c,** Location of somatic piRNA clusters with *flam*-like properties in (a) *D. mojavensis*, (b) *D. virilis*, and (c) *D. ananassae*. **d-g,** Examples of candidate unistrand cluster syntenic to the *D. ananassae* candidate (c) on the 5' end (near a *cac*-orthologous gene). Genomic LTR content across 100 kb windows is shown. Predicted LTR content (blue, plus strand; red, minus strand) and total repeat content (grey) is shown across the whole genome (100 kb windows). Cluster loci are indicated at the top and selected contig/chromosome names are indicated at the bottom.

Pipeline overview

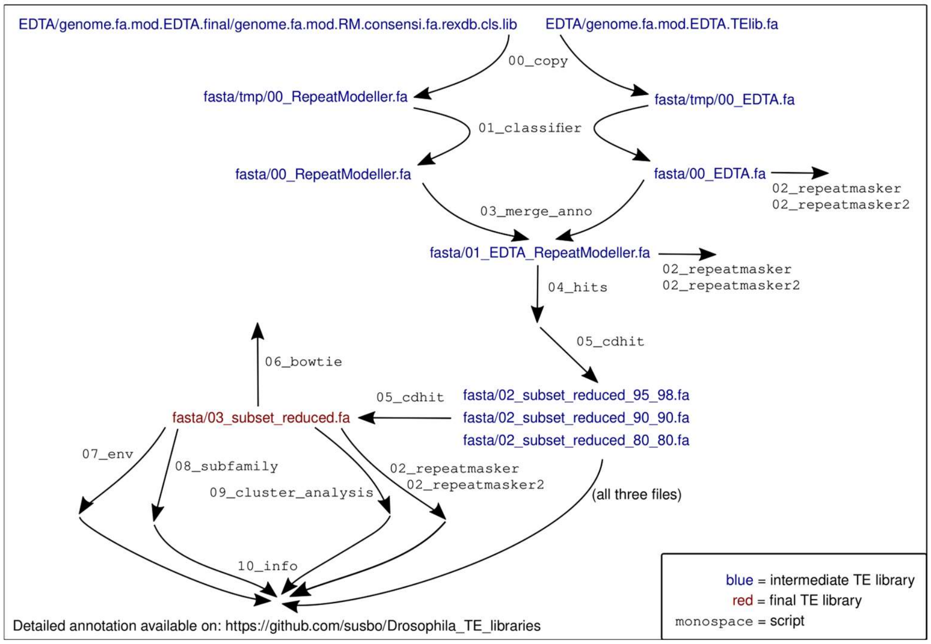


**Fig. S17: Overview of *de novo* transposon consensus sequence construction**

Illustration of how the raw TE libraries into processed to obtain a set of curated TE consensus sequences (see "Transposon_libraries" at https://github.com/susbo/Drosophila_unistrand_clusters). Arrows labelled with mono-space font indicate bash scripts and blue/red text indicate some of the key output files.

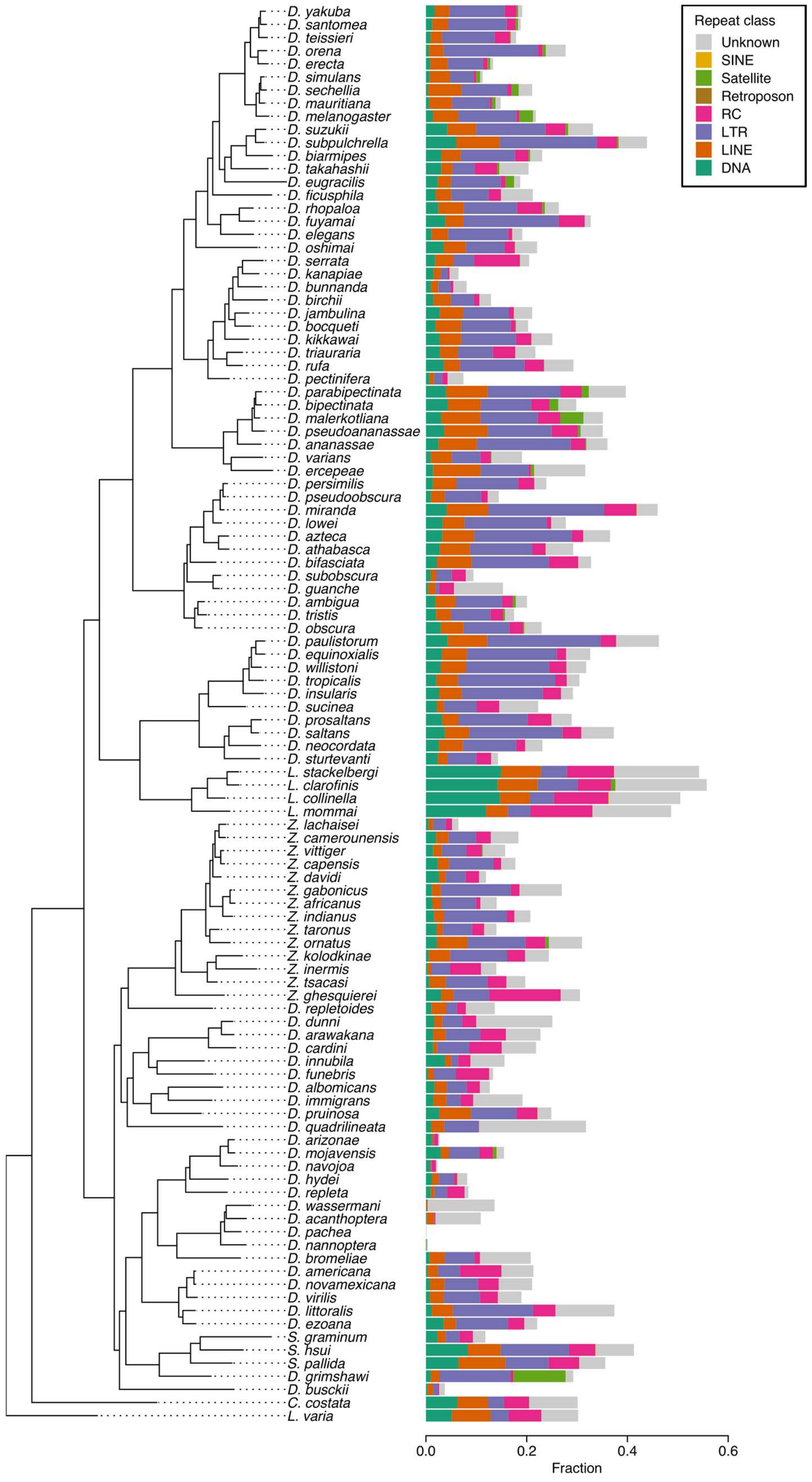


**Fig. S18: Repeat content of *Drosophila* species**

Genomic interspersed repeat content per species predicted using EDTA.

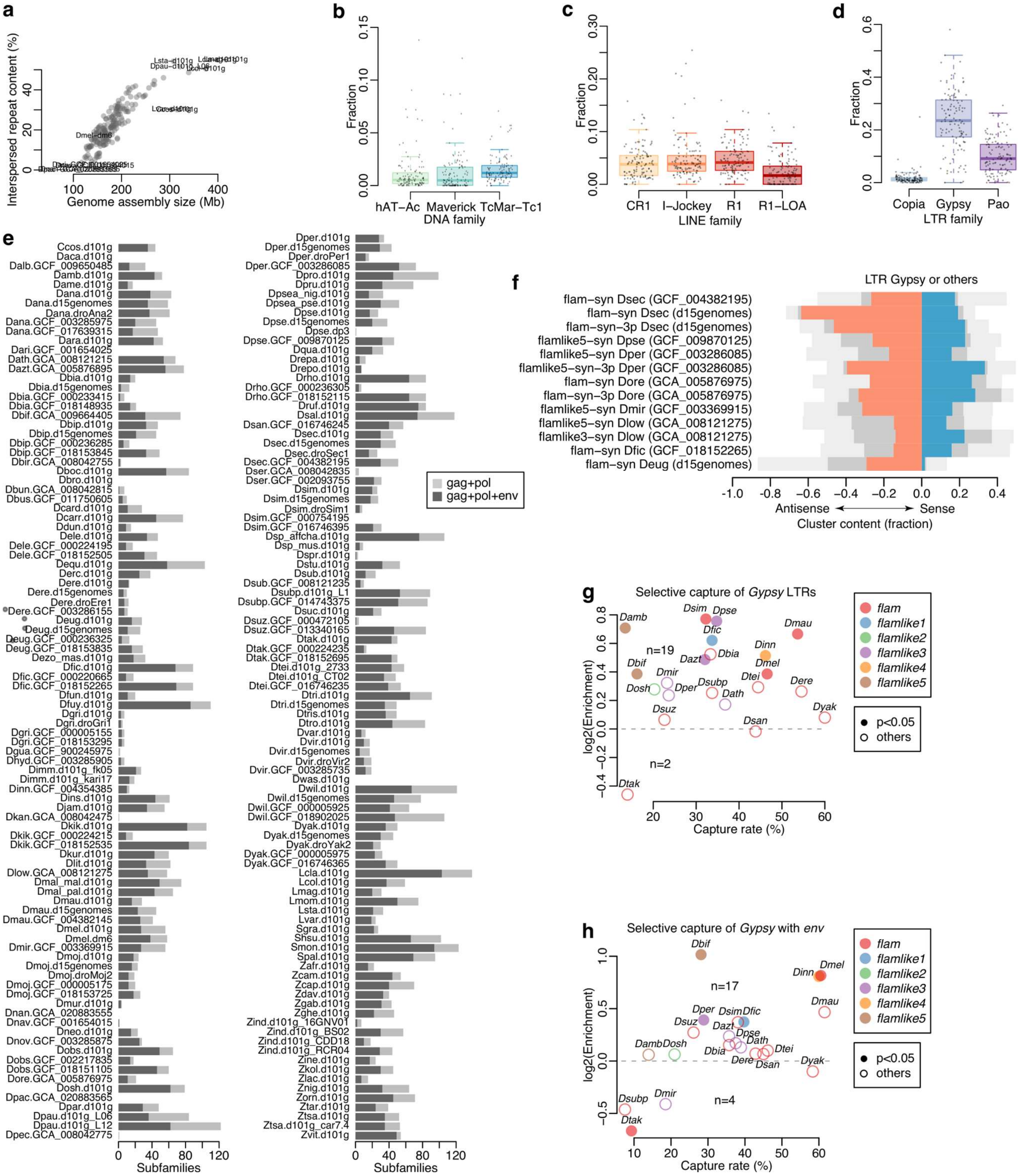


**Fig. S19: *De novo* TE libraries enable detailed cluster analyses**

**a**, Interspersed repeat content correlates with genome assembly size. Each data point represents one genome assembly (n=193). Selected assemblies are indicated. **b**, Boxplot showing fraction of interspersed repeat content for the most abundant DNA families (mean above 1%). Each data point represents one species (n=119). Species with multiple genome assemblies are represented by their mean. **c**, Same as in (b) but showing LINE families. **d**, Same as in (b) but showing LTR families. **e**, Bar plots showing the number of high-confidence (*gag* and *pol* present) LTR *Gypsy* subfamilies identified per genome assembly. Subfamilies are separated based on the presence (red) or absence (grey) of *env*. Each data point represents one genome assembly (n=193). **f**, Bar plot showing LTR contribution (left, antisense; right, sense) to total transposon content across all annotated *flam*-syntenic clusters. *Gypsy* elements are shown in red (antisense) or blue (sense) and other LTR elements are shown in grey. Total transposon composition is shown in light grey. **g**, Scatterplot showing *Gypsy* capture rate per cluster against *Gypsy* enrichment (*Gypsy* against all LTRs calculated for the indicated cluster against the rest of the genome). Only LTRs with both *gag* and *pol* having at least one good genomic hit were included in the analysis. One-sided enrichment p-values were calculated using bootstrap (n=1000 replicates). **h**, Scatterplot showing *Gypsy env* capture rate per cluster against *Gypsy env* enrichment (*Gypsy* with *env* against all *Gypsy* calculated for the indicated cluster against the rest of the genome). Only LTRs with both *gag* and *pol* having at least one good genomic hit were included in the analysis. One-sided enrichment p-values were calculated using bootstrap (n=1000 replicates). Boxplots show median (central line), interquartile range (IQR, box), and minimum and maximum values (whiskers, at most 1.5*IQR).

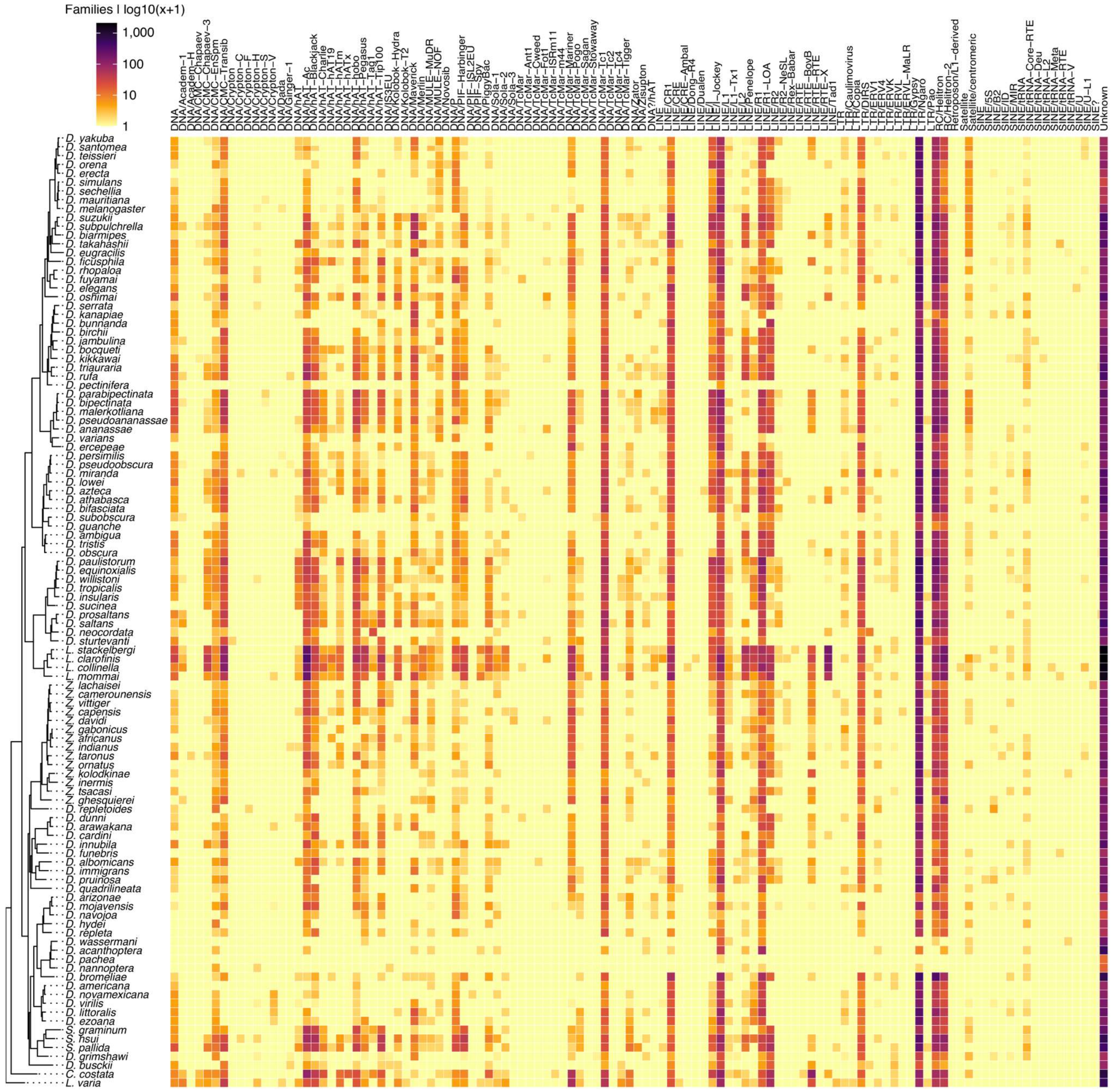

**Fig. S20: Overview of identified subfamilies per TE family across *Drosophila* species**

Identified number of subfamilies per TE families across indicated *Drosophila* species. Only families detected in at least one species are shown. Detailed numbers are available in Table S4.

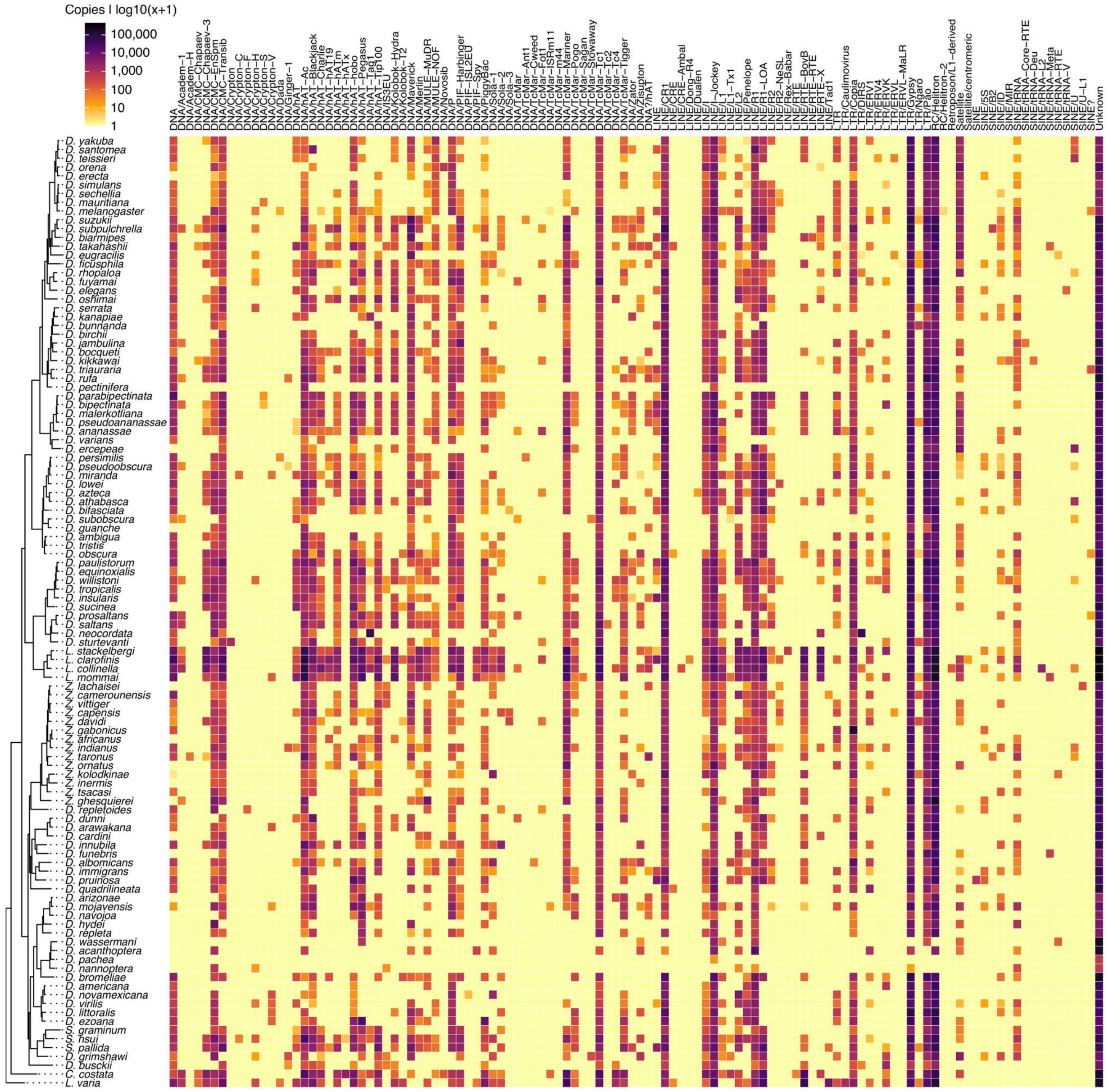

**Fig. S21: Overview of genomic copies per TE family across *Drosophila* species**
Identified number of genomic copies for TE families across indicated *Drosophila* species. Only families detected in at least one species are shown. Detailed numbers are available in Table S5.

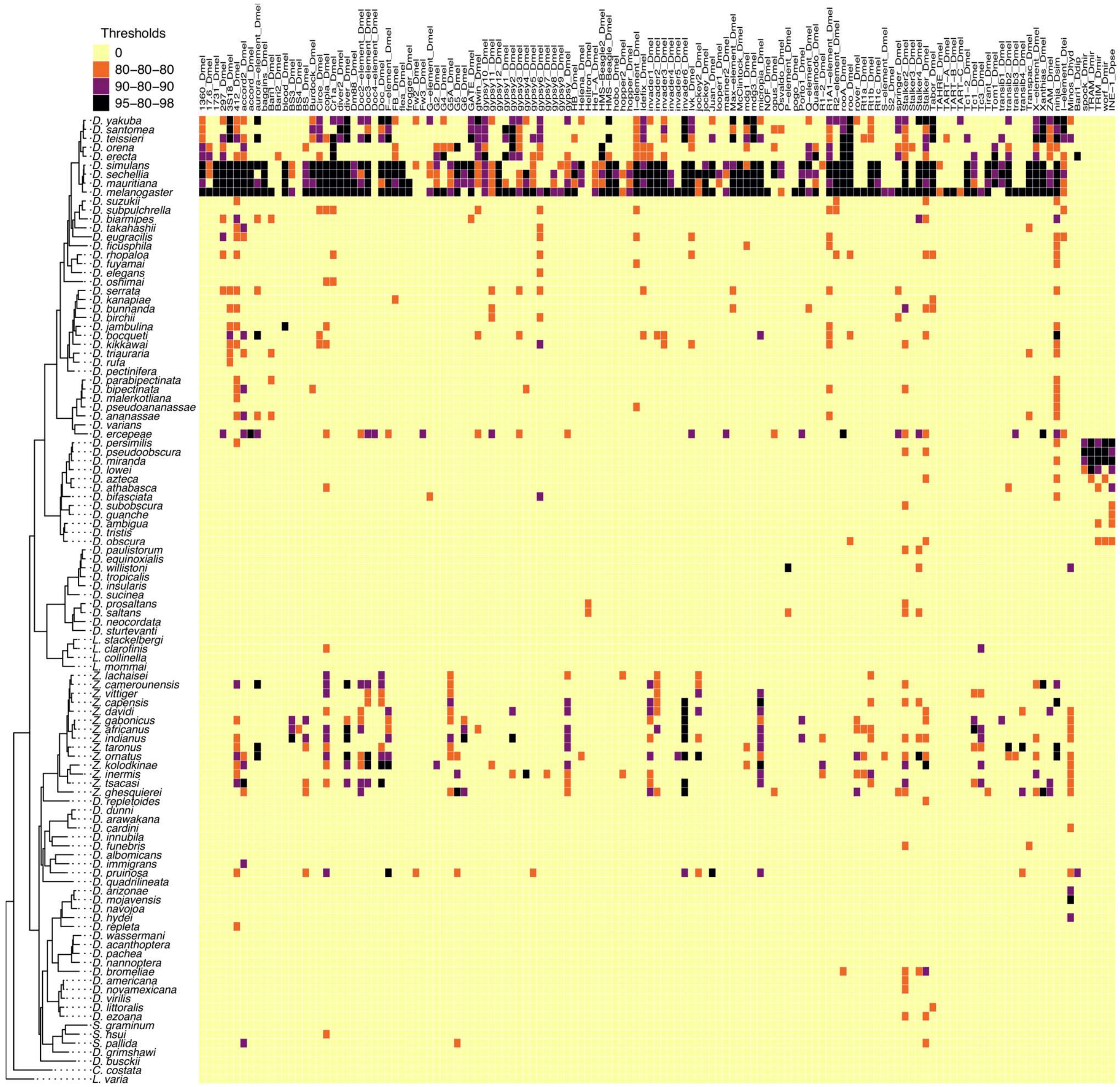


**Fig. S22: Overview of previously described TE subfamilies across *Drosophila* species**
Representation of TE subfamily identification (n=155) across *Drosophila* species. A TE was considered to belong to a certain subfamily with very high confidence (black) if it had a hit with at least 95% identity, across 98% of the query, and at least covering 80 nt (i.e., 95-80-98 threshold). Similarly, medium-confidence (purple, 90-80-90) or low-confidence (orange, 80-80-80) hits are indicated. This data is also available in Table S6.

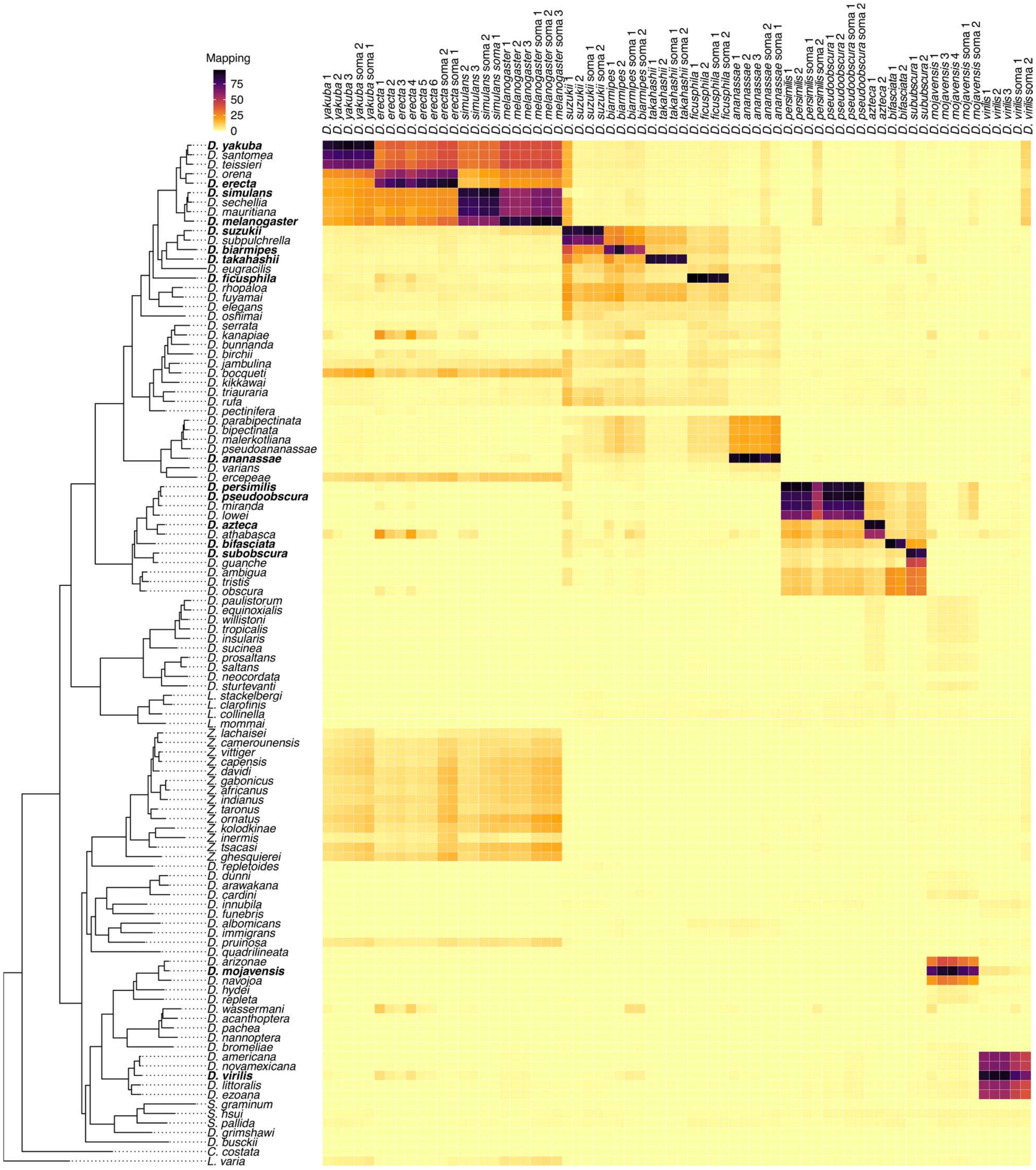

**Fig. S23: Alignment of sRNA-seq across *Drosophila* species**

Alignment of sRNA-seq across indicated *Drosophila* species. Species that were sequenced are indicated in bold. Horizontal lines such as the one for *D. ercepeae* indicate potentially shared piRNAs due to horizontal transmission of TEs. Vertical lines such as the one for *D. persimilis* indicate likely index swapping with *melanogaster* subgroup species sequenced at the same lane. See also Fig. S24 for a similar analysis using RNA-seq. Abbreviations: FC, follicle cells.

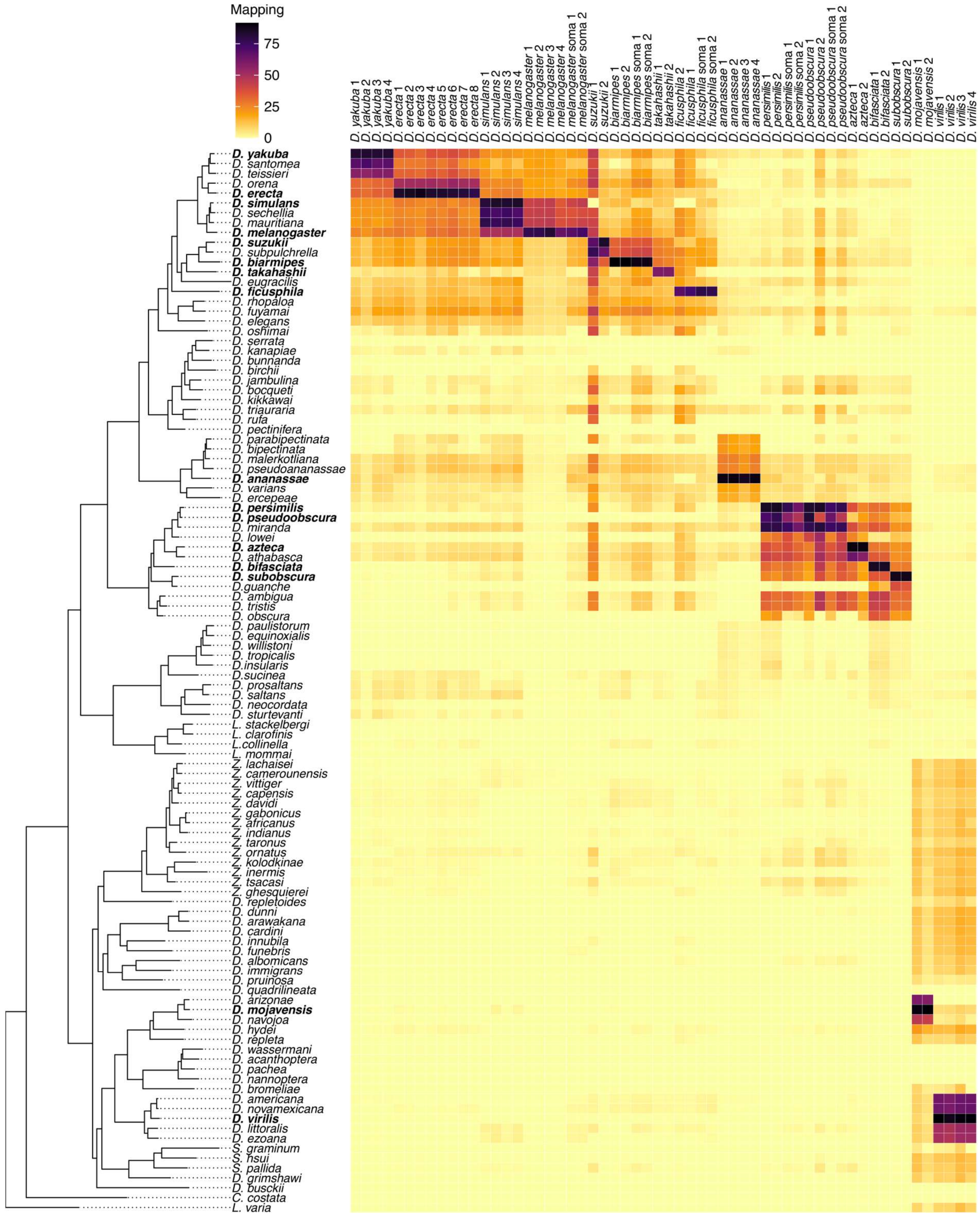


**Fig. S24: Alignment of RNA-seq across *Drosophila* species**

Alignment of total RNA-seq across indicated *Drosophila* species. Species that were sequenced are highlighted in bold.

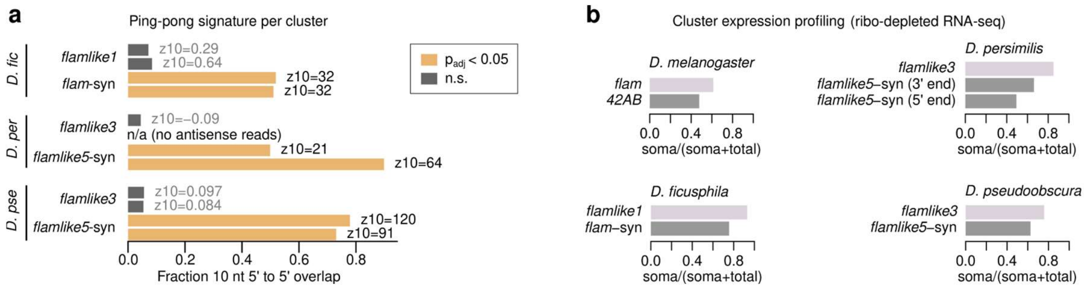


**Fig. S25: *Flam*-like clusters display somatic expression and reduced ping-pong signature**

**a,** Barplots showing ping-pong signature across somatic *flam*-like and germline control clusters in *D. ficusphila*, *D. persimilis*, and *D. pseudoobscura* (2 biological replicates each). Ping-pong signature was quantified as the fraction of 10 nt overlaps piRNAs mapping to opposite strands and assessed using a two-tailed *z*-test with Bonferroni correction. **b**, Barplots showing relative cluster expression in soma-enriched (soma) and whole ovary (total) RNA-seq libraries. Signal was first converted to cpm to correct for differences in sequencing depth. Pooled counts from 2-4 biological replicates.

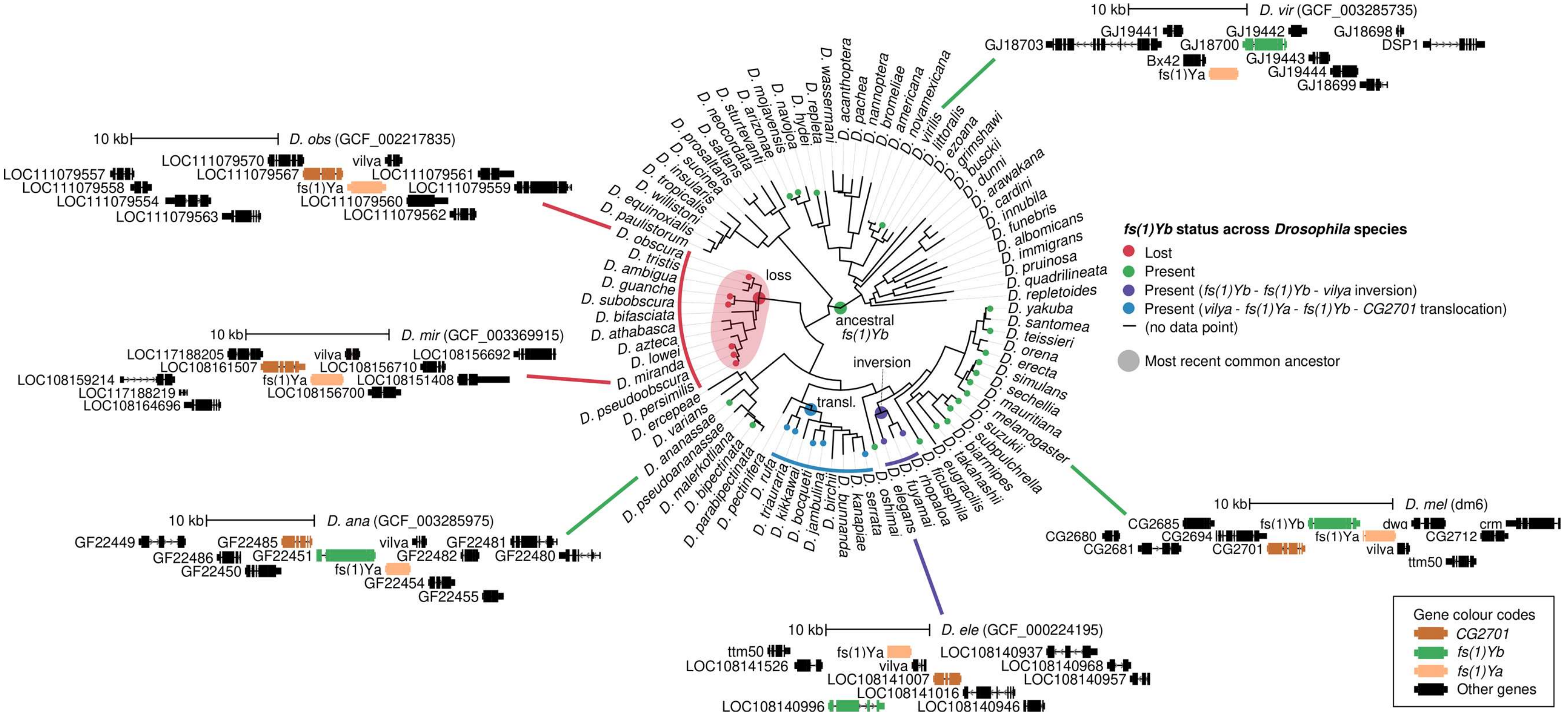


**Fig. S26: Conservation of *fs(1)Yb* across *Drosophila* species**

Phylogenetic tree summarising *fs(1)Yb* status across *Drosophila* species. Only genome assemblies with NCBI gene predictions were evaluated. Syntenic regions were identified in each genome and annotated to reflect whether *fs(1)Yb* was present (green) or absent (red) at the expected location next to *CG2701* and/or *fs(1)Ya*. Additionally, the presence of *fs(1)Yb* together with specific inversions (purple) or translocations (blue) are indicated. Selected examples of gene synteny or lack thereof are shown surrounding.
